# Non-invasive, Real-time Detection of Vascular Disorders in Mice using Bright SWIR-emitting Gold Nanoclusters and Monte Carlo Image Analysis

**DOI:** 10.1101/2020.01.31.928382

**Authors:** Zhixi Yu, Benjamin Musnier, Maxime Henry, K. David Wegner, Benoit Chovelon, Agnès Desroches-Castan, Arnold Fertin, Ute Resch-Genger, Sabine Bailly, Jean-luc Coll, Yves Usson, Véronique Josserand, Xavier Le Guével

**Author notes:** E-mail:* &.

## Abstract

We present here a new approach for non-invasive high resolution whole-body vascular imaging in depth by combining water-soluble and bright SWIR-emitting gold nanoclusters revealing an anisotropic surface charge with Monte Carlo image processing of the images. We applied and validated this approach to quantify vessel complexity in transgenic mice presenting vascular disorders.

---

*In vivo* infrared imaging has experienced major breakthroughs over the past few years with potential applications in cancer and cardio-vascular diagnostics^1^. Hongjie Dai’s team was among the first who developed emitters for the shortwave infrared region (SWIR, 900-1700 nm), also called NIR II. Due to the weak photon absorption, the low autofluorescence, and reduced scattering by tissues at these wavelengths compared to NIR I (700-900 nm) and the visible region, they were able to reach a high spatial and temporal resolution through a few millimeters of tissue sufficient to image brain blood circulation through the intact skull^2^ and in ischemic femoral arteries^3^. Using SWIR imaging, Bawendi *et al.* recently evaluated the metabolic turnover rates of lipoproteins in several organs in real-time as well as the heartbeat and breathing rates in awake and unrestrained animals^4^. Based on these data, they generated a detailed three-dimensional quantitative flow map of the mouse brain vasculature, which demonstrates the high potential of SWIR-imaging applications.

However, only few SWIR emitting contrast agents are available that possess high quantum yields (QYs), good biocompatibility and low accumulation in organs. Although *in vivo* imaging studies have been performed using SWIR-emitting quantum dots (QDs)^4^, the down-converted emission of lanthanide-based nanomaterials^5^, and new organic donor– acceptor–donor (D-A-D) type organic fluorophores^6^, these materials still have some drawbacks. This includes for example, their possible toxicity related to the toxicity of their constituents in the case of most QDs and low QYs < 1 % for the organic molecules. A relatively new class of NIR II emitters are ultra-small gold nanoclusters (Au NCs). Recent studies with Au NCs stabilized by zwitterionic sulfobetaine ligands (AuZw) showing a high QY (3.8%), enabled to detect blood vessels using photoluminescence (PL) and revealed efficient renal clearance^7^.

In this work, we developed a new type of water-soluble Au NCs with emission in the SWIR, which exhibit a high brightness, a high photostability, long blood circulation times, and a low toxicity. We demonstrate the ability of this promising new contrast agent to image the vascular network of mice with vascular disorders in the second window of the SWIR spectrum (1250-1700 nm) with a more than one-fold enhancement in spatial resolution enabled by the use of Monte Carlo image processing. The non-invasive imaging processing, segmentation and analyzes were validated in transgenic mice inactivated for the Growth factor *Bmp9* that were previously described with vascular disorders^8–9^ based on the measurement of the fractal dimension of vessels.

The here developed Au NCs, named AuMHA/TDT, were prepared by a wet chemistry route using mercaptohexanoic acid (MHA) and tetra(ethyleneglycol) dithiol (TDT) as co-ligands with the molar ratio Au: ligand =1:4 and MHA: TDT= 1:3 (**Figure 1a**, *see SI for the experimental details*). Mass spectrometry revealed a high monodispersity of an 11 kDa species (Figure **S1**). The average size of the semi-crystalline metal core was determined by high resolution transmission electron microscopy (HR-TEM) to 2.1±0.6 nm (**Figures 1b, S2**) and the hydrodynamic diameter of AuMHA/TDT in water to 1.90±0.02 nm as derived from DOSY-NMR (**Figure S3**). The water-soluble AuMHA/TDT NCs have a negative surface charge at pH 7 of around ζ= −20 mV, which renders them very stable over the long period of time. The addition of the short dithiol molecules TDT on the Au NC surface results in a striking modification of the optical properties, as recently reported by us for another type of anisotropic Au NCs prepared with hexa(ethylene glycol) dithiol as co-ligand^10^. Indeed, the incorporation of TDT leads to the presence of new NIR absorbance features at 800 nm, 910 nm and 1140 nm (**Figure 1c**). The anisotropic surface of Au NCs also influences drastically the PL signal in the SWIR region with a 9-fold increase of PL intensity in water accompanied by PL emission at longer wavelengths (**Figure 1d**). The spectral position and shape of the emission is independent of the excitation wavelength (**Figure S4**) and the average lifetime derived from the multiexponential decay detected at λ_em_.= 930±20 nm is <τ_int_.>= 451.9 ns (**Figure S5**). We compared the PL of AuMHA/TDT to the emission of other water-soluble Au NCs synthesized in our laboratories, i.e., AuMHA_10_, AuZw_11_, and Au_25_SG_1812_ in different spectral windows in the SWIR. The images in **Figure 1e** show an up to 12-fold more intense signal for AuMHA/TDT using long pass filters above 1250 nm confirming the superior brightness of our new Au NCs in water as compared to the existing SWIR-emissive ones. The PL features of AuMHA/TDT remain unchanged in water and in the presence of 10% serum (**Figure S6**). Illumination studies in water, with NaCl (0.9%), and in the presence of bovine serum albumin (BSA;50 mg/mL) at different incubation times demonstrate the high photostability of these emitters (**Figure S7**). Surprisingly, in the presence of blood, the PL increases over time. A similar behavior has also been reported for ICG^13^ and is tentatively ascribed to specific interactions with proteins/complement components or uptake by red blood cells.

**Figure 1.**
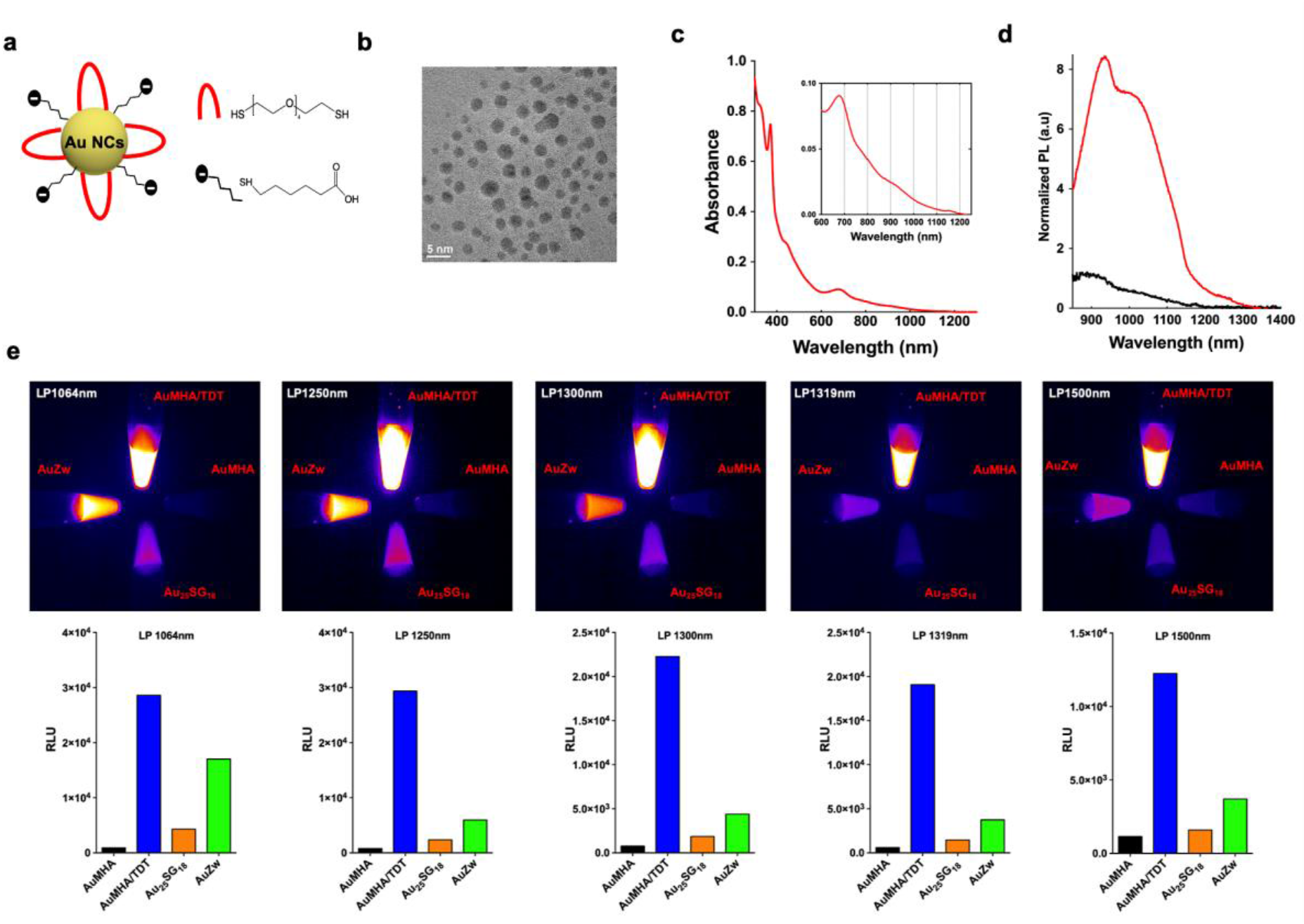
**a**. Scheme of the Au NCs AuMHA/TDT. **b.** HR-TEM images of AuMHA/TDT. **c.** Absorbance spectra of AuMHA/TDT. **d.** PL spectra of AuMHA (black line) and AuMHA/TDT (red line) (λ_exc_. 830 nm). **e.** SWIR PL of AuMHA, AuMHA/TDT, Au_25_SG_18_, and AuZw (300 μM in water) under NIR excitation (λ_exc_. 830 nm) using LP1064 nm (1 ms), LP1250 nm (5 ms), LP1300 nm (10 ms), LP1319 nm (25 ms), and LP1500 nm (250 ms).

Cytotoxicity experiments (not shown) have been conducted using MTT assays with a human embryonic cell line (HEK 293) and human (U87, Colo 829) or murine (4T1) cancer cell lines in the presence of increasing concentrations of AuMHA/TDT up to 90 μM (~1 mg/mL). The absence of significant cell death suggests the low toxicity of these contrast agents and allowed us to move forward to *in vivo* studies.

The half-life of Au NCs in mice is highly dependent on the hydrodynamic diameter of the NCs ^11, 14^ and their density^15^. For instance, small Au_25_SG_18_ NCs show an extremely fast clearance with a half-life of less than 2 min as compared to 6 min for the larger AuZw_11_. By elemental analysis (ICP-MS) after AuMHA/TDT administration, we measured a half-life of t_1/2α_= 0.43±0.05 h and a half-elimination of t_1/2β_ = 19.54 ±0.05 h in nude mice (**Figure 2a**). These two values are more than 5 times longer than those obtained with the previous applied Au NCs. This prolonged blood circulation time enables the detection of SWIR signals from AuMHA/TDT in the blood vessels in mice up to 30 minutes after injection.

**Figure 2.**
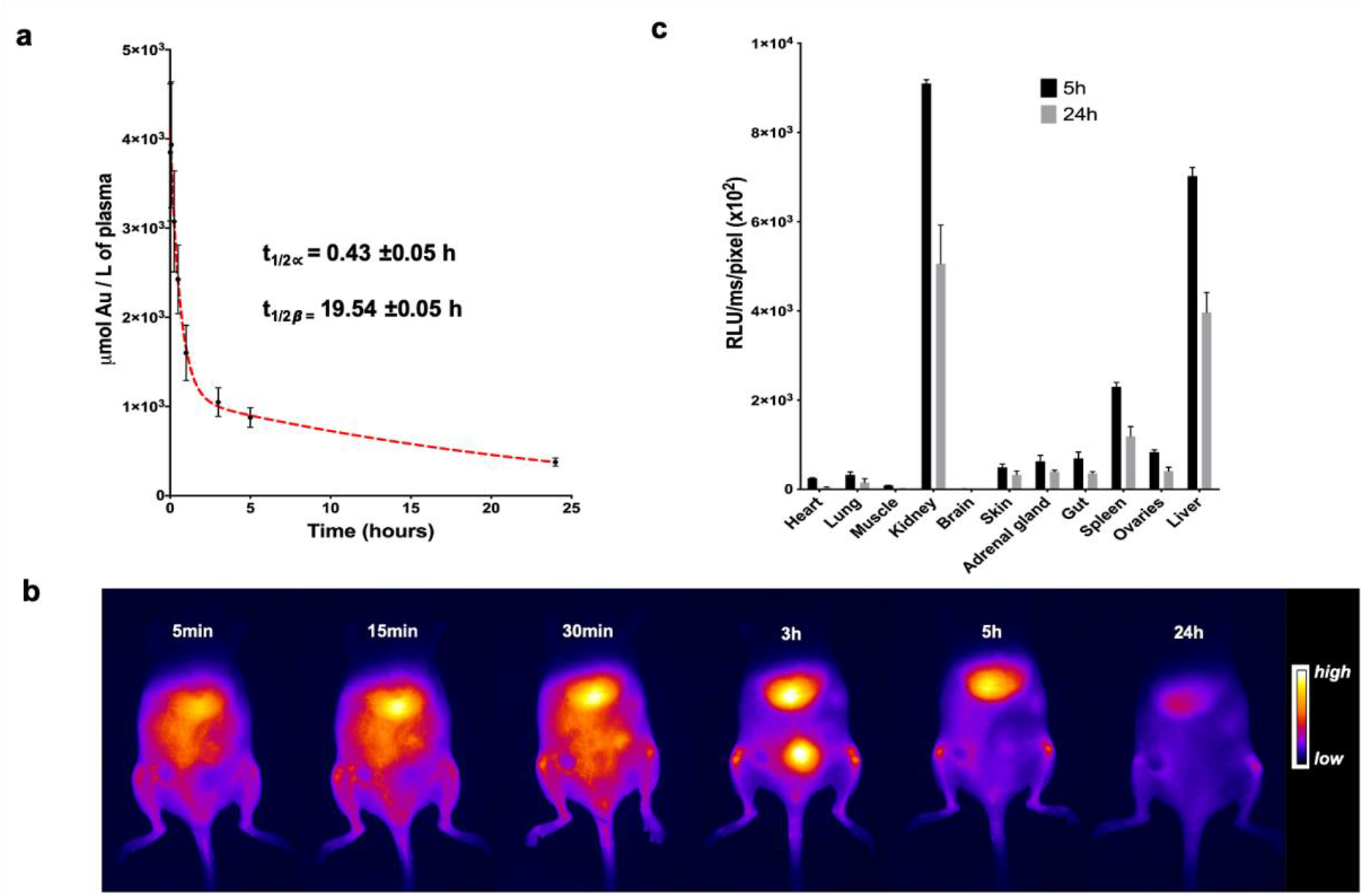
Biodistribution and pharmacokinetics of AuMHA/TDT after intravenous injection in nude mice (360 μM; 200 μL per mouse). **a.** *In vivo* pharmacokinetics determined by ICP-MS measurements on plasma samples taken at different time points (n=4 mice per time points). **b.** *In vivo* pharmacokinetic by fluorescence in mice over 24h (λ_exc_. 830 nm; LP1250 nm). **c.** *Ex vivo* 2D fluorescence signal of AuMHA/TDT in isolated organs 5h and 24h post injection (n=3 mice per time points).

*In vivo* fluorescence measurements also show the renal clearance of these ultra-small particles with signal in the bladder at 3h post injection and with an elimination from the liver observed between 5h to 24h, respectively (**Figure 2b**).

One of the major advantages of the SWIR spectral window is the low autofluorescence, which enables to reduce the amount of the contrast agent necessary to achieve a high signal to noise ratio. In our case, we could detect AuMHA/TDT NCs in *in vitro* samples at concentrations as low as 80 nM with a signal-to-noise ratio of 5.5 (**Figure S8**). This enabled us to follow the biodistribution of AuMHA/TDT in mice by fluorescence imaging after 5 h or 24 h post-injection.

Results from *ex vivo* fluorescence measurements in different organs (**Figure 2b**), confirmed by ICP-MS measurements (**Figure S9**), suggest a renal elimination of the Au NCs and an accumulation of the Au NCs mainly in the kidney, the liver, and the spleen with PL signal dropping by 40% between 5 h and 24 h, which suggests an elimination or a metabolization of AuMHA/TDT over time.

Non-invasive real-time SWIR imaging was performed after intravenous injection of AuMHA/TDT (200 μL; 360 μM) *in vivo* in 129/Ola mice using a 830 nm laser as light excitation source (50 mW/cm^2^) and a long-pass filter LP1250 nm in the emission channel. Under these conditions, the AuMHA/TDT NCs provided a clear and outstanding visualization of the vascular network as shown in **movie 1**.

As expected, there is a clear improvement of the spatial resolution in the SWIR is strongly improved as compared to the NIR I (**Figure S10**). Despite this very good *in vivo* spatial resolution, we nevertheless performed post-image processing with the Monte Carlo Constrained Restoration (MCR) method to further improve the spatial resolution in depth and to overcome the scattering from the skin and the tissues. Image processing greatly enhances the spatial resolution as seen in **Figure 3a**. Comparing the quality of the images before and after MCR using a quantitative measure of the contrast (**Figure 3b**) demonstrates an improvement in contrast by one order of magnitude, highlighting the potential of this restoration. The determination of the transverse section of blood vessels from the SWIR images of a mouse leg depicted in **Figure 3c** highlights the excellent resolution after MCR processing and the ability to obtain highly detailed *in vivo* imaging of the whole-body blood vasculature and of the blood brain vasculature (**Figure S11**).

**Figure 3.**
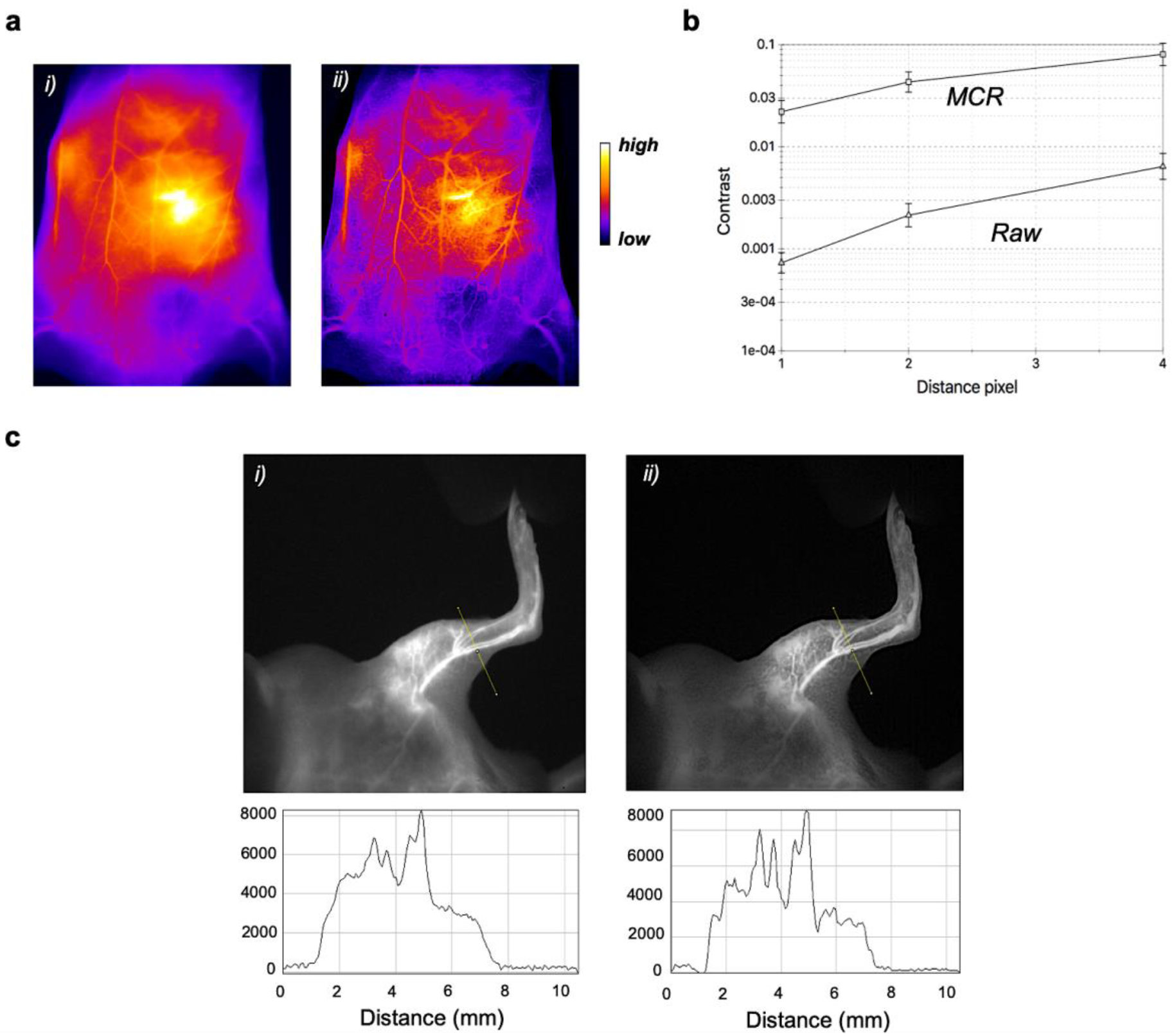
SWIR imaging of blood vasculature in WT 129/Ola mice after intravenous injection of AuMHA/TDT. **a.** Non-invasive imaging of the mouse ventral area (i) before and (ii) after MCR processing (false colors). **b.** Quantification of contrast enhancement per pixel with and without MCR processing on the whole ventral area. **c.** Images of the left leg of the mouse (i) before and (ii) after MCR processing with their respective intensity profile across a line of interest drawn in the inset images above.

Image processing was then pushed one step further using a high-pass filter to reduce scattering deeper under the skin. Restorations of the first movie by MCR plus the high pass filter underline the striking improvement of the resolution (**movie 2**) allowing the visualization of the vascular network at more than 4 mm penetration depth (**Figure 4a**). We could also easily track the contrast agent (SWIR-emitting AuMHA/TDT) in mice especially in the first seconds after injection (**movie 3, Figure 4b**).

**Figure 4.**
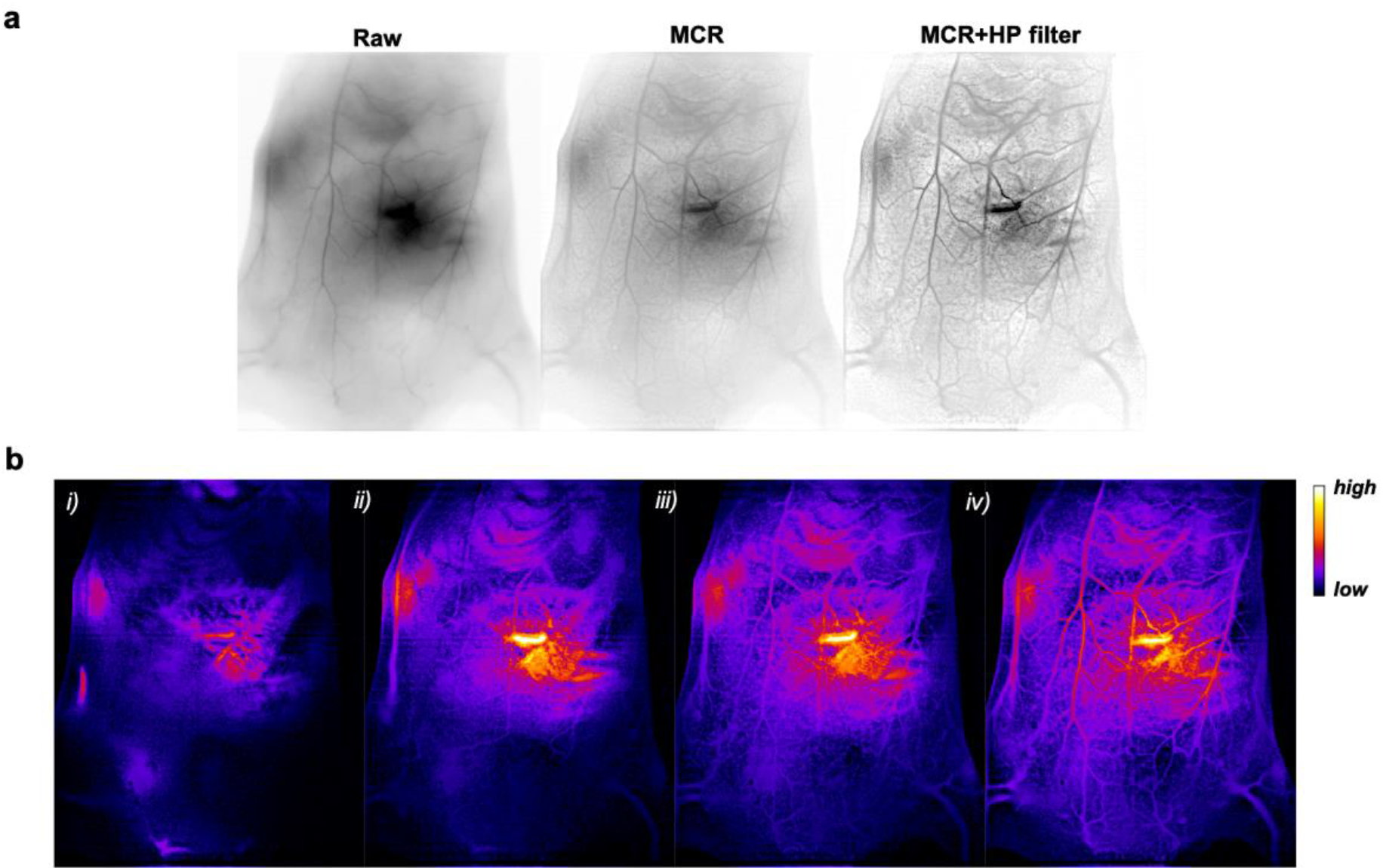
**a.** *In vivo* SWIR imaging (reverse contrast) of WT 129/Ola mice vasculature before imaging processing (raw) and after Monte Carlo constrained restoration (MCR) and an additional filtering (MCR+HP filter). **b.** MCR+HP filter treated SWIR images (false colors) i) 1.5s, ii) 5s, iii) 25s, and iv) 65s after AuMHA/TDT bolus intravenous injection (360 μM; 200 μL).

To demonstrate the potential of combining SWIR imaging, image restoration, and image analysis for biomedical applications, non-invasive vascular imaging was performed in mice previously described with vascular disorders due to the inactivation of the gene encoding *Bmp9*^8–9^ and compared to wild-type mice (WT). *Bmp9* has been identified as a high affinity ligand for the endothelial specific receptor ALK1 (activin receptor-like kinase 1), that is mutated in the rare vascular disease HHT (hereditary Haemorrhagic Telangiectasia)^16^. We have recently shown that *Bmp9* deletion in the 129/Ola genetic background results in dilated liver vessels that ultimately lead to liver fibrosis^9^.

*Bmp9-*KO mice and WT mice were placed in supine position and were injected intravenously with AuMHA/TDT for real-time SWIR imaging from t=0 to t=15 min. MCR was performed on raw images followed by Frangi’s filtering and segmentation (*see methods in SI*) as illustrated **in Figure 5a**. Then, analyses of the segmentation, which provides blood network mapping were plotted considering the length of the vessels or the number of vessel “branches” (segments delimited by forks or crossings) as a function of the fractal dimension (**Figure 5b**). Fractal dimensional is referred here as the level of complexity of vessels with a numerical value between 1 (straight line tube) to 2 (highly swaying vessel in 2-dimensional section). Analyses show a clear distinction in skin vessels between *Bmp9*-KO and WT mice considering both vessel lengths and branches. The significant 4% increase of the fractal dimension for Bmp9-KO compared to WT mice, which is correlated to the vessel distortion, confirms then the hypothesis of tortuous vessels due to a defect in vessel maturation^9^. Thus, this non-invasive method allowed us for the first time to detect a vascular disorder present in the *Bmp9*-KO mice with such a high level of precision.

**Figure 5.**
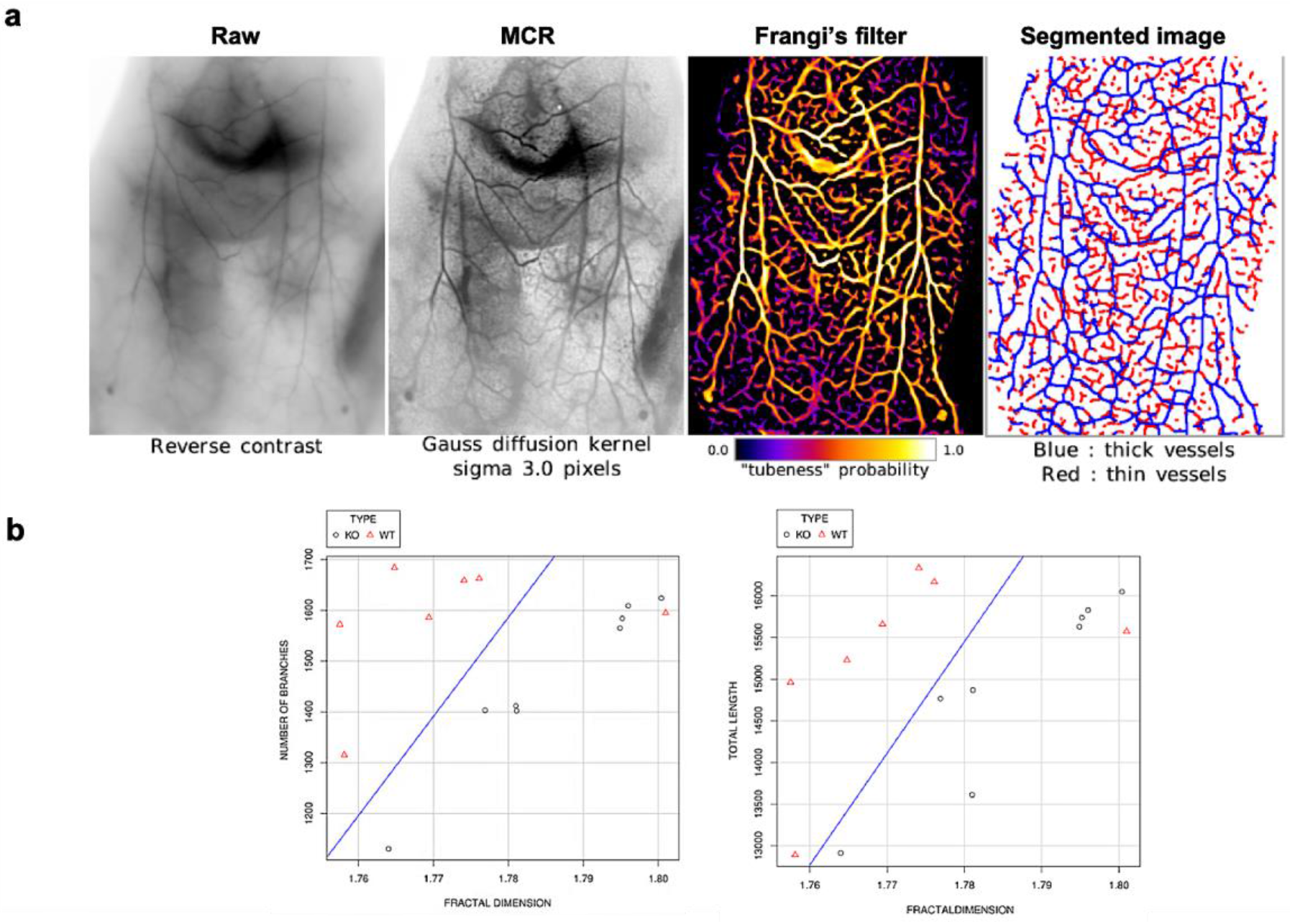
**a.** SWIR images of a *Bmp9*-KO mouse after MCR processing, Frangi’s filter, and segmentation. **b.** Statistical analyses of blood vessel length and branches as a function of fraction dimension performed on *Bmp9*-KO mice (n= 4) and WT mice (n= 4). The blue line highlights the discrimination between the two groups.

In conclusion, we report here the design of a new bright SWIR-emitting contrast agent with prolonged circulation and efficient elimination that enables the visualization of the vascular network with enhanced-spatial resolution in depth using a series of MCR processing steps. Both image processing and segmentation analyses enabled to distinguish non-invasively vascular disorders in mice with good confidence. To the best of our knowledge, this is the first time that such results were obtained non-invasively in depth and in real time using whole-body optical imaging. This study also highlights the considerable advantages of combined research on new contrast agents and on image processing and analyses to improve the sensitivity of SWIR imaging for advanced biomedical applications.

## Supporting Information

Supporting Information is available from Springer or from the author.

## Acknowledgements

XLG would like to thank Cancéropôle Lyon Auvergne Rhône-Alpes (CLARA), Plan Cancer (C18038CS) and ARC (R17157CC) for their financial support. XLG would like to thank Ines Häusler for the electron microscopy images (The TEM images were carried out as part of the DFG core facility project “Berlin Electron Microscopy Network (Berlin EM Network)”) and Muriel Jourdan for the NMR measurements. KDW acknowledges the European Union’s Horizon 2020 research and innovation programme under the Marie Sklodowska-Curie grant agreement No.846764. This research was funded by the Institut National de la Santé et de la Recherche Médicale (INSERM, U1036), the Commissariat à l’Energie Atomique et aux Energies Alternatives (CEA, DRF/IRIG), the University Grenoble Alpes (UGA, BCI), the Fondation pour la Recherche Médicale (FRM),

---

Ultra-small gold particles are used as SWIR contrast agent to detect a vascular disorder with a high level of precision non-invasively in depth and in real time using whole-body optical imaging.

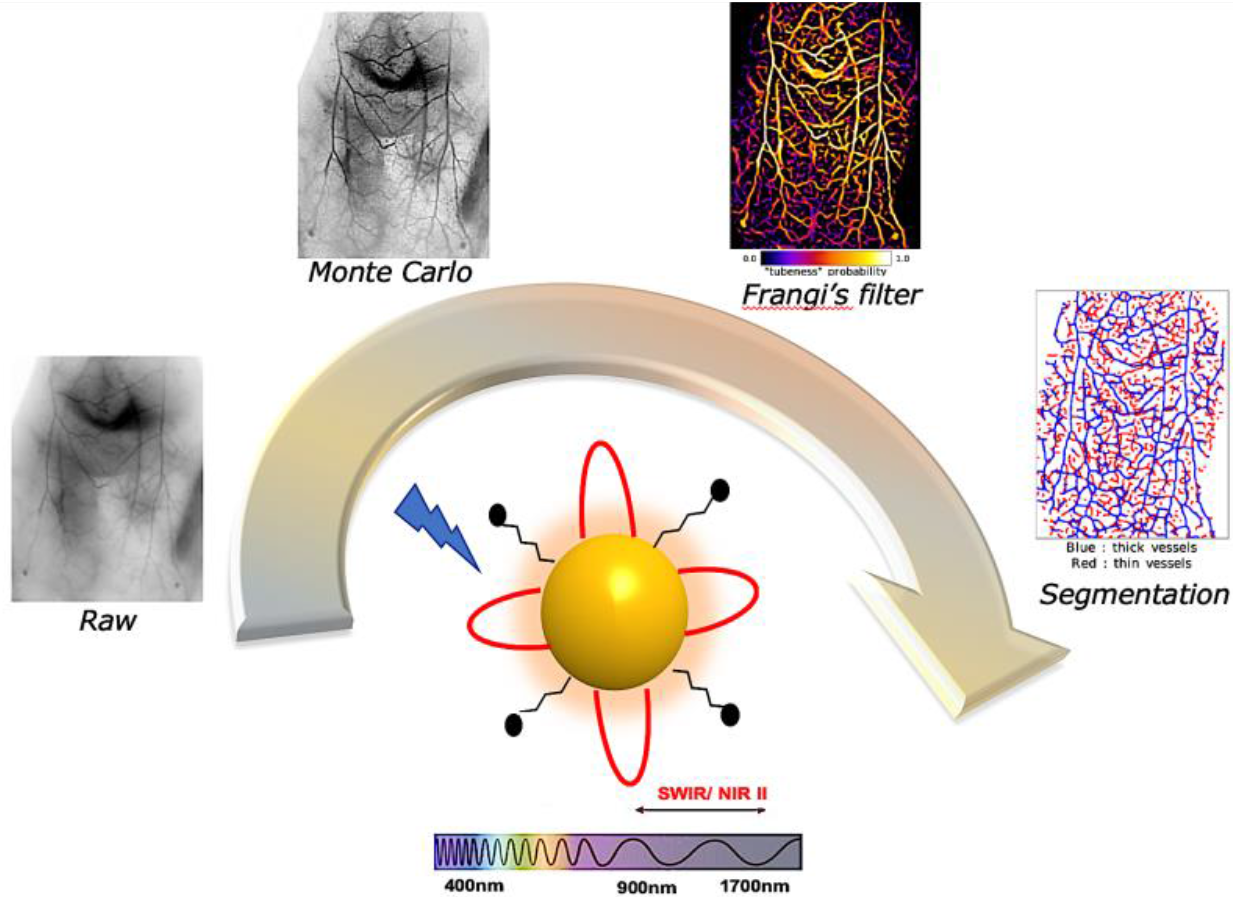

