## Supplementary material for "Non-invasive, Real-time Detection of Vascular Disorders in Mice using Bright SWIR-emitting Gold Nanoclusters and Monte Carlo Image Analysis": SI

| | |
|---|---|
| Methods & instrumentations | 2 |
| Synthesis Au NCs | 7 |
| Physico-chemical characterisations | 8 |
| Photo-physical characterisations | 9 |
| *In vivo* studies | 12 |

### 1- Methods & instrumentations

- **Matrix Assisted Laser Desorption Ionisation - Time of Flight (MALDI-TOF).** Au NCs sample was diluted in the matrix CHCA (Alpha-Cyano-4-Hydroxy-3-Iodocinnamic Acid) with 0.1% trifluoroacetic acid (TFA) in a mixture water/acetonitrile 50/50 v/v. Measurements were performed in positive linear mode on an Autoflex speed instrument from Brucker.
- **Nuclear magnetic resonance spectroscopy (NMR)** experiments were carried out at 298 K with a Bruker AVANCE III 500 MHz spectrometer equipped with a cryo-probe Prodigy. For each sample the concentration was ~2 mM in $D_2O$ and pH 7. Diffusion ordered NMR spectroscopy (DOSY) experiments were run using the standard “ledbpg2s” Bruker sequence with linear gradient stepped between 2% and 98%. 32 scans were recorded for each gradient step. Data processing were performed using the maximum entropy algorithm from Dynamics Center, a Bruker’s NMR software to obtain the diffusion coefficient D. An average value of D was used for the hydrodynamic diameter (HD) calculation according to the Stokes-Einstein equation which assumes that molecules are spherical:

$$HD = kBT/ 3D\pi\eta$$

where kB is the Boltzmann constant, T is the temperature, η is the viscosity of the solvent (ηD2O = 1.232 10-3 Pa.s at 298K). Standard 1D and 2D (NOESY, COSY, TOCSY and HSQC) spectra were recorded using standard presaturation on the water signal.

- **Zeta potential** of Au NCs dispersed in water or in PBS buffer with 10% serum were measured with a Zetasizer from Malvern instruments.
- Metal core sizes were determined by **high resolution transmission electron microscopy** with an 200 kV monochromated TEM using dispersed Au NCs on ultra-fine carbon films.
- Absorption spectra of diluted AuNC samples were recorded on a **UV-vis-NIR spectrophotometer** Cary5000 between 350 and 1700 nm.

- **Steady-state photoluminescence** spectra were measured from 600 – 1750 nm with a calibrated FSP 920 (Edinburgh Instruments, Edinburgh, United Kingdom) spectrofluorometer equipped with a nitrogen-cooled PMT R5509P.
- **Time-resolved measurements** were performed in the wavelength region of 930±20 nm using a FLS 920 (Edinburgh Instruments, Edinburgh, United Kingdom) lifetime spectrofluorometer equipped with an EPL-510 (Edinburgh Instruments, Edinburgh, United Kingdom) picosecond pulsed diode laser (excitation wavelength of 510±10 nm; power of 5mW) and a fast PMT R2658P from Hamamatsu, respectively.
- **Relative measurements of photoluminescence** QYs (Φf,x) were performed using the dye IR125 dissolved in dimethylsulfoxide (DMSO as reference. The QY of this dye was previously determined absolutely to Φf,st = 0.23). The relative QY were calculated according to the formula of Demas and Crosby6, see equation below.

$$\Phi_{f,x} = \Phi_{f,st}\frac{F_x}{F_{st}}\cdot\frac{f_{st}(\lambda_{ex,st})}{f_x(\lambda_{ex,x})}\cdot\frac{n_x^2(\lambda_{ex,x})}{n_{st}^2(\lambda_{ex,st})}$$

The subscripts x, st, and ex denote sample, standard, and excitation respectively. f(λ_ex ) is the absorption factor, F the integrated spectral fluorescence photon flux, and n the refractive index of the solvents used (DMSO for IR125; water for Au nanoclusters).

All spectroscopic measurements were done in a 1 cm quartz cuvettes from Hellma GmbH at room temperature using air-saturated solutions.

- **NIR I imaging** was performed with a NIR 2D-Fluorescence Reflectance Imaging device (Fluobeam 800®, Fluoptics, France). The excitation is provided by a class 1 expanded laser source at 780 nm and the irradiance on the imaging field is 10 mW/cm2. The fluorescence signal is collected by a CCD through a High pass filter with a high transmittance for wavelength > 830 nm.
- **SWIR imaging** was performed using a Princeton camera 640ST (900-1700 nm) coupled with a laser excitation source at λ= 830 nm (50 mW/cm2). We use short-pass excitation filter at 1000 nm

(Thorlabs) and long-pass filters on the SWIR camera from Semrock (LP1064 nm, LP1319 nm) and Thorlabs (LP1250 nm, LP1300 nm, LP1500 nm). 25 mm or 50 mm lenses with 1.4 aperture (Navitar) were used to focus on the samples or mice.

Tubes containing AuNC solution and 10 µL drops of each samples were placed in front of the camera using the 50 mm lens and various long pass filters (LP1064 nm, LP1250 nm, LP 1300 nm, LP1319 nm, LP1500 nm). Analyses were performed using FIJI software.

Mice were imaged after intravenous injection (200 µL of Au NCs at 360 µM) using the 25 mm or 50 mm lenses and LP1250 nm at different exposure times (25ms to 2s). Ex vivo fluorescence imaging on isolated organs and plasma samples were performed using the 50 mm lens and LP1064 nm.

- **Inductively coupled plasma-mass spectrometry (ICP-MS)** was performed to determine Au content in organs and in plasma samples at different time point using a Thermo X serie II, spectrometer (Thermo Electron, Bremen, Germany), which was equipped with an impact bead spray chamber and a standard nebulizer (1 $mL.min^{-1}$). For sample preparation, the organs and plasma samples were weighted before addition of nitric acid (final concentration 1%) and Au content was determined using an external linear calibration curve (between 10 and 100 µg/L of Au(III)). Indium was used as the internal standard. Determinations were carried out in triplicate.

- ***In vivo* experiments**

For the biodistribution study, six weeks old NMRI female nude mice (Janvier, France) were anesthetized (air/isoflurane 4% for induction and 1.5% thereafter) and were injected intravenously via the tail vein with 200 µL of Au NCs at 360 µM. In vivo SWIR fluorescence imaging was performed before and 5 and 24 hours after injection. Mice were euthanized at 5 or 24 hours post injection (n = 3 mice per time point) and organs were harvested for ex vivo fluorescence imaging and ICP measurements.

For the pharmacokinetic study, three other mice were injected and blood samples were collected before and 1, 5, 15, 30, 60, 180, 300, and 1440 min after injection and were centrifuged (10 min at 2000 g) to separate plasma. Plasma pharmacokinetic was obtained from fluorescence imaging and ICP-MS measurements after analyses through a non-compartment model (GraphPad Prism 7.00, GraphPad Software, La Jolla California USA).

*Noninvasive SWIR imaging on mice with vascular disorders.* Generation of *Bmp9*-KO mice in the 129-P2/Ola-Hsd genetic background (named here for easiness 129/Ola) has been previously described[1-2]. Briefly, *Bmp9*-KO mice in the C57BL/6 genetic background were obtained from Dr Se-Jin Lee (Johns Hopkins University, Baltimore, MD) and back-crossed for 10 generations with 129-P2/Ola-Hsd wild-type mice (Harlan France, Gannat, France).

All animals experiments followed the institutional guidelines formulated by the European Community for the Use of Experimental Animals were approved by ethics committees (CEA ethic committee for animal breeding and Cometh38 for in vivo imaging) and the French Ministry of Research and Education. (agreement APAFIS#9436-2017032916298306 and APAFIS#21916-2019082710189095_v4).

**- Image restoration of SWIR fluorescent images**

SWIR already provides stunning fluorescent images in vivo, however these images still suffer from light diffusion by the tissues and the quality of such images can still be improved using image restoration techniques. We used a Monte Carlo constrained reconstruction (MCR) algorithm based on an original idea by Frieden *et al.*[3] for restoring binary images then extended to fluorescent image deconvolution by Colicchio *et al.*[4]. This algorithm has the advantage of perfectly preserving the amount of information of the image (intensity integral) and achieve powerful contrast and resolution enhancements, while minimizing the ringing artefacts usually encountered with regular iterative constrained deconvolution. However, Monte Carlo algorithms are very computer intensive and parallelisation schemes must be used to speed up calculations.

- **Assessment of image enhancement**

The enhancement of fluorescent images was both assessed by visual inspection (qualitative assessment) and by using a quantitative measurement. This was achieved by quantifying the contrast $C_d$ within the image at different neighbourhoods (distance $d = 1, 2$ and $4$ pixels). The contrast $C_d$ was expressed as the gradient integral of the image.

$$C_d = \frac{\sum_1^{r-d}\sum_1^{c-d}\left(\frac{\delta i}{\delta_d x}^2 + \frac{\delta i}{\delta_d y}^2\right)}{\langle i\rangle^2}$$

with

$$\frac{\delta i}{\delta_d x} = i(x+d,y) - i(x,y)\ ;\quad \frac{\delta i}{\delta_d y} = i(x,y+d) - i(x,y);\ \ \langle i\rangle = \frac{\sum_1^r\sum_1^c i(x,y)}{r\cdot c}$$

where *i(x,y)* is the intensity at pixel *(x,y)* ; *d* is the extent of the partial derivate (in pixels) ; $\delta i/\delta_d x$ the partial derivate of intensity along *x* and $\delta i/\delta_d y$ the partial derivate of intensity along *y* at pixel *(x,y)*; *r* is the number of rows and *c* the number of columns in the image.

- **Vessel detection and analysis**

The analysis of the vascularization was performed by image analysis of the restored images. We used a classical Frangi's filter that was designed to enhance tubular structures in a grey level image[5]. The advantage of Frangi's filter is a good immunity to noise that reduces over-detection. Frangi's filter provides a "tubeness" or "vesselness" probability image to which a probability threshold can be applied in order to obtain a binary mask of the vascular network. After, iterative thinning of the binary mask, a skeleton is obtained and a neighbourhood analysis is applied in order to extract branches, forks and crossings. The statistical analysis of the various features extracted from the skeleton such as: the fractal dimension, total length, the number of branches, the number of forks & crossings; is used to quantify possible differences between mouse strains.

#### 2- Synthesis of Au NCs

Chemical products were purchased in Sigma-Aldrich (France) and deionized water was used for all synthesis.

We slightly modified a protocol described by Musnier et al.[6] to produce the SWIR-emitted Au NCs using the initial molar ratio Au:Ligand = 1:4. Briefly 250 µL of $HAuCl_4.3H_20$ (20 mM) was added to 4.8 mL water followed by 4 mL of the thiolated ligand mixture mercaptohexanoic acid (MHA, 5 mM)) / tetra(ethyleneglycol) dithiol (TDT, 5 mM) changing colour from yellowish to slightly pale cloudy with a volume ratio MHA/TDT= 3mL/1mL. After 5 min, 250 µL of NaOH (1M) was added dropwise leading to almost colourless sols. After 5 min, 150 µL $NaBH_4$ (20 mM in 0.2 M NaOH) was introduced dropwise under mild stirring and kept under stirring at 350 rpm for 8 hours. Purification of the **AuMHA/TDT** on 3 kDa cut-off filter column (Amicon) were repeated 3 times to stop the reaction and sols were kept stored in the fridge before characterization.

The Au NCs, Au MHA[6], AuZwMe[27] and $Au_{25}SG_{18}$[8] were synthesized following the protocols described in the literature.

### 3- Structural characterization

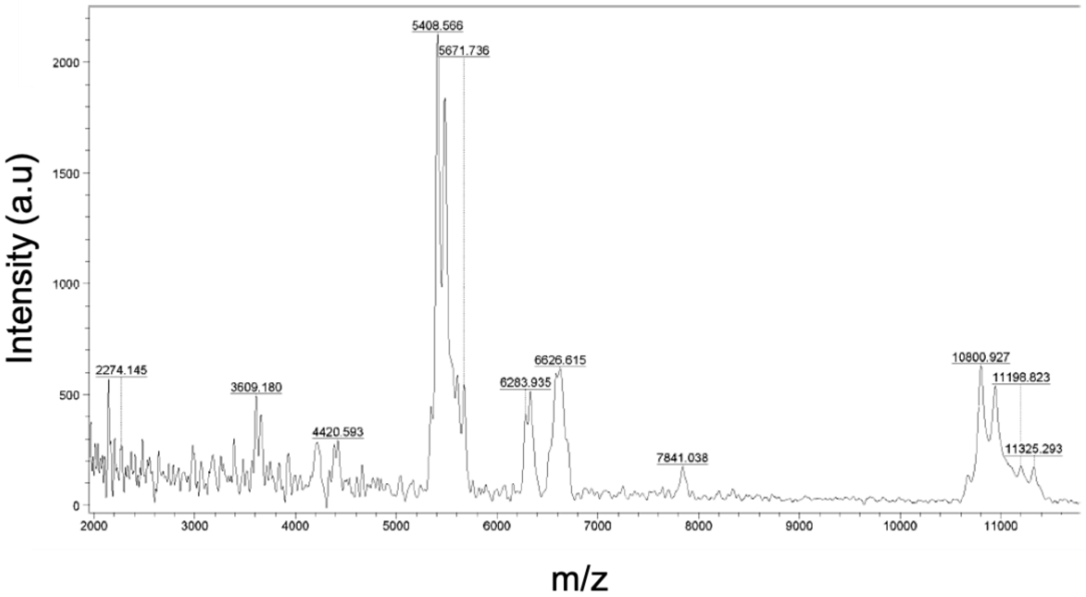


Figure S1. MALDI-TOF pattern of AuMHA/TDT in linear positive mode.

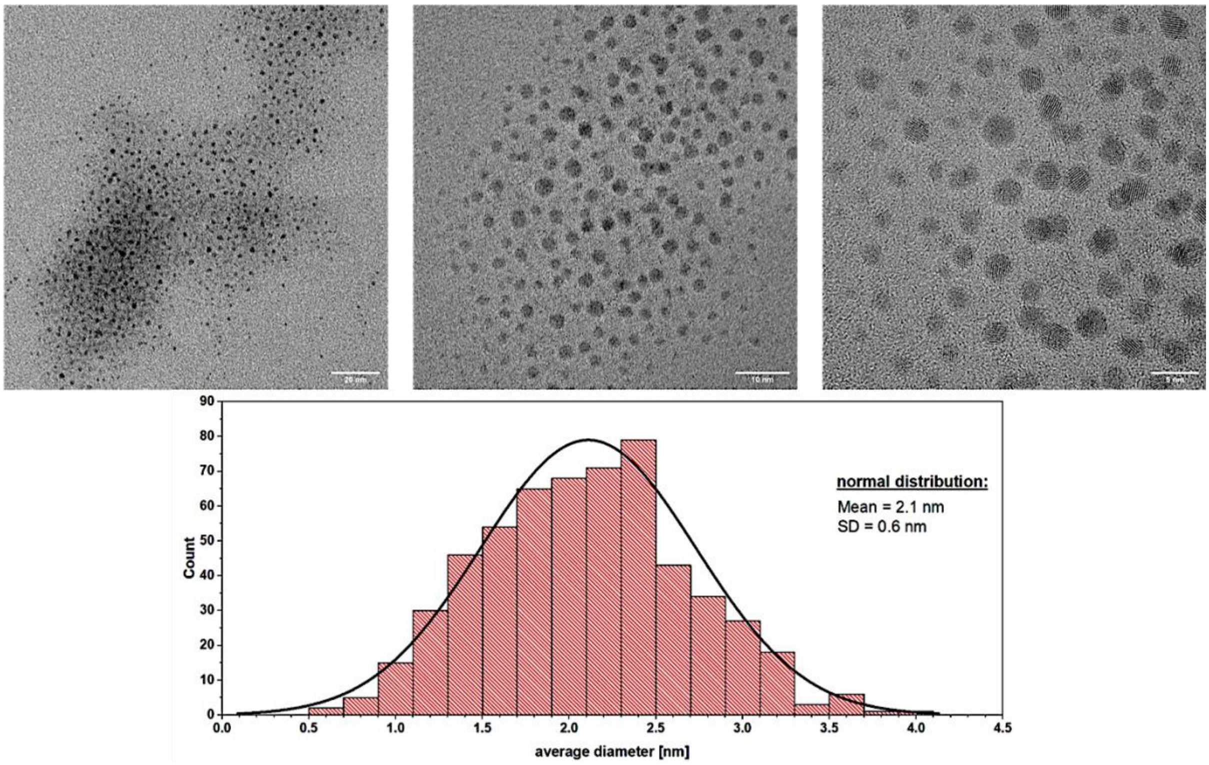


Figure S2. High-resolution TEM of AuMHA/TDT sample and statistical analysis of particle size based on more than 100 particles.

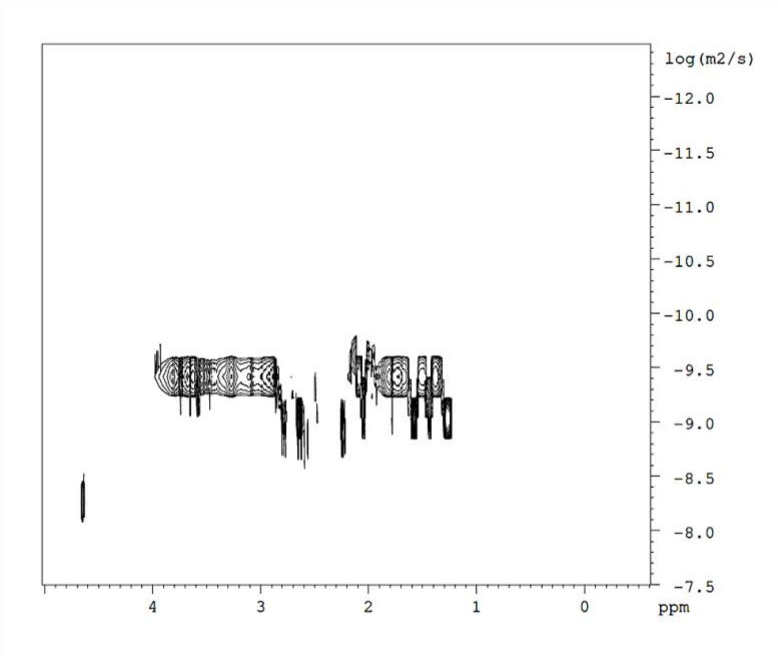


Figure S3. DOSY-NMR of AuMHA/TDT in $D_2O$. Coefficient diffusion D = 1.8 $10^{-10}$ $m^2/s$, which corresponds to a hydrodynamic diameter $\varnothing_{HD}$ = 1.90±0.02 nm.

### 4- Photophysical characterization

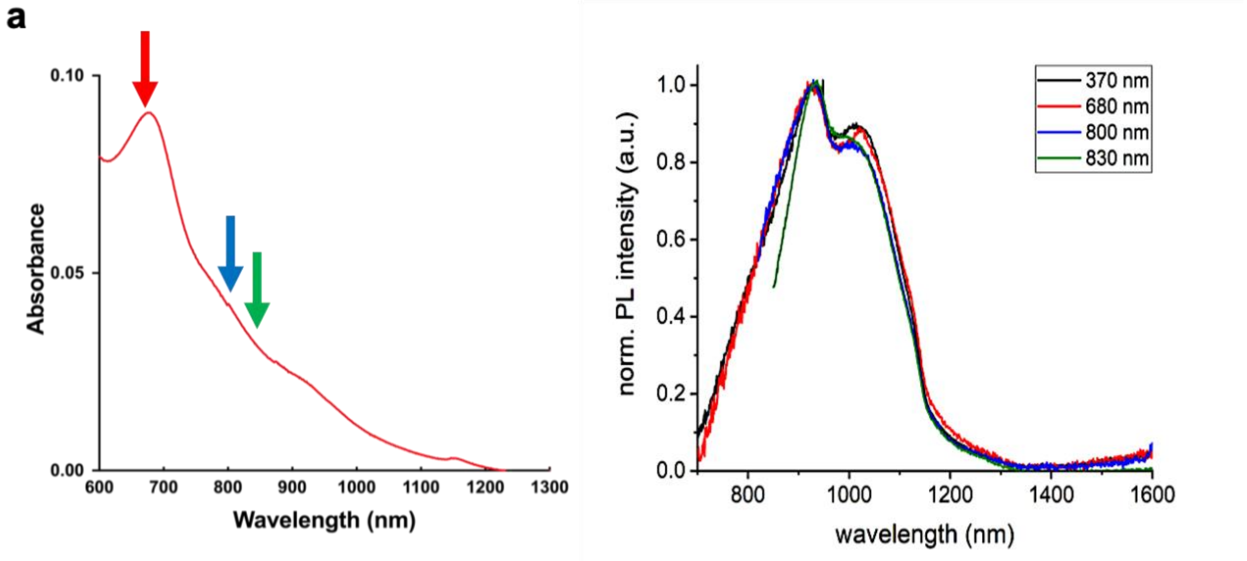


Figure S4. (a) Absorption spectrum of AuMHA/TDT in water and (b) normalized PL spectra using different excitation wavelengths.

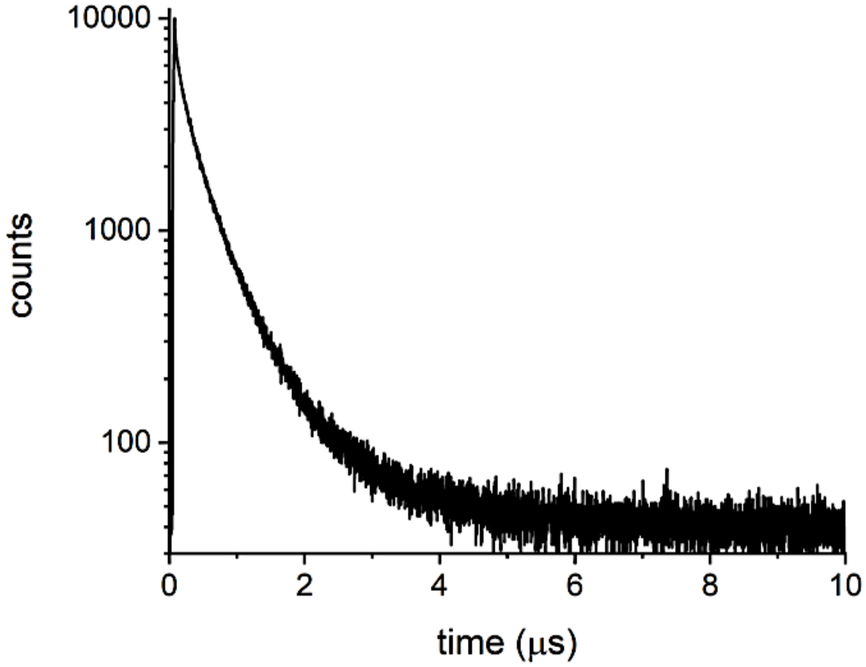


| $F_1$ (%) | $\tau_1$ (ns) | $F_2$ (%) | $\tau_2$(ns) | $F_3$ (%) | $\tau_3$ (ns) | $F_4$ (%) | $\tau_4$ (ns) | <τ> int. |
|---|---|---|---|---|---|---|---|---|
| 4.2 | 1.5 | 8.0 | 63.4 | 56.6 | 326.7 | 31.2 | 838.8 | 449.0 |

Figure S5. Fluorescence lifetime measurement of AuMHA/TDT in water.

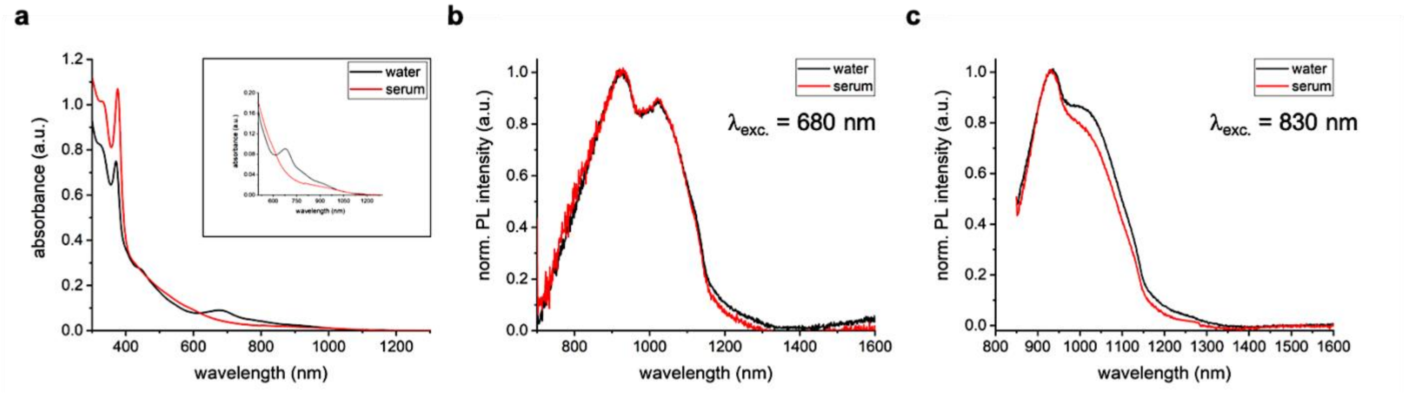


Figure S6. (a) Absorbance and (b,c) fluorescence spectra of AuMHA/TDT dispersed in water and in serum (10% calf-serum in DMEM).

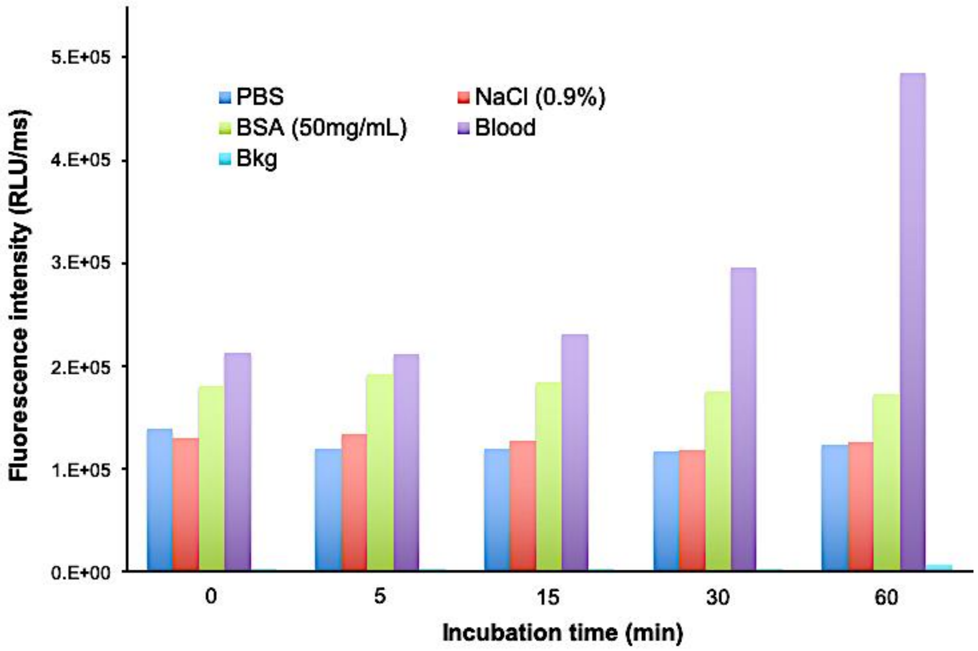


Figure S7. Fluorescence intensity of AuMHA/TDT incubated at 37°C in different media for 0, 5, 15, 30, and 60 min measured by SWIR camera (LP1064 nm) and the measured background (Bkg).

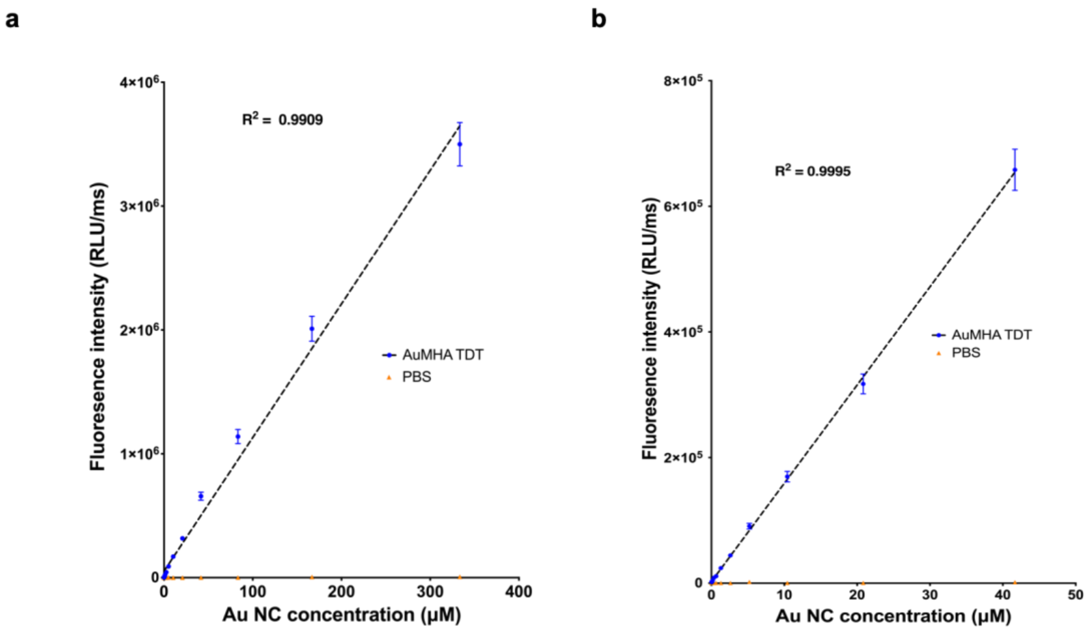


Figure S8. *In vitro* AuMHA/TDT fluorescence signal detection (blue dot) in PBS between 0 and 360 μM (a) and between 0 and 50 μM (b) by SWIR imaging (LP1064 nm).

### 5- *In vivo* studies

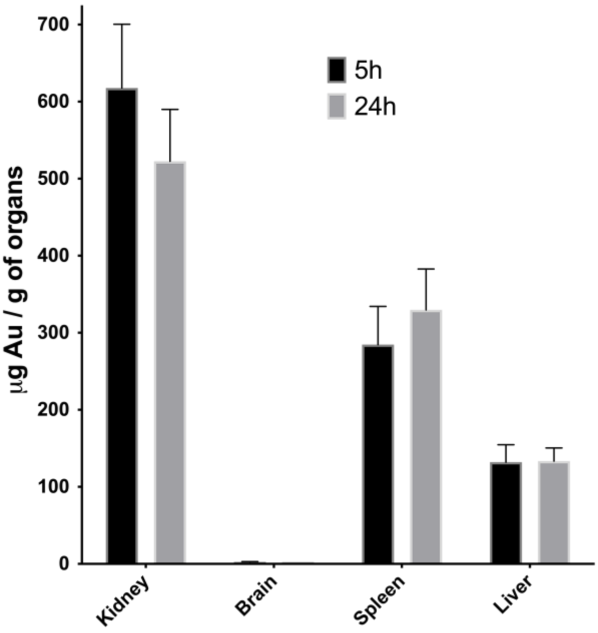


Figure S9. ICP-MS measurement of gold in organs (n=3) taken from mice 5 and 24h after *i.v* injection of AuMHA/TDT (200μL; 360μM).

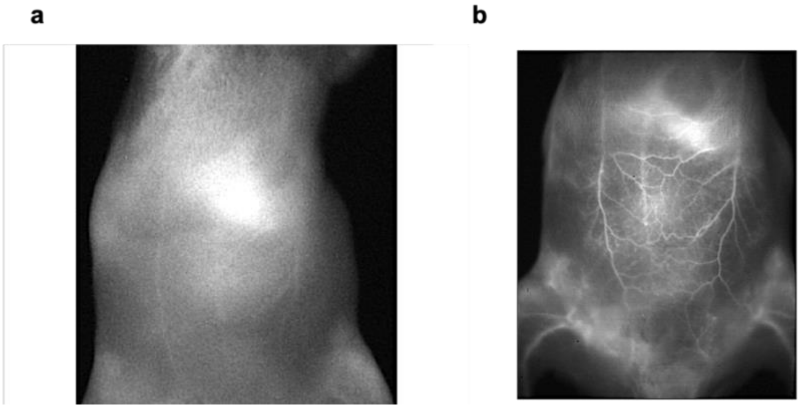


Figure S10. (a) NIR I (ex 780 nm; Em > 830 nm) and (b) SWIR imaging (LP1250 nm) of mice 15 min after *i.v* injection of AuMHA/TDT (200 µL at 360 µM in PBS).

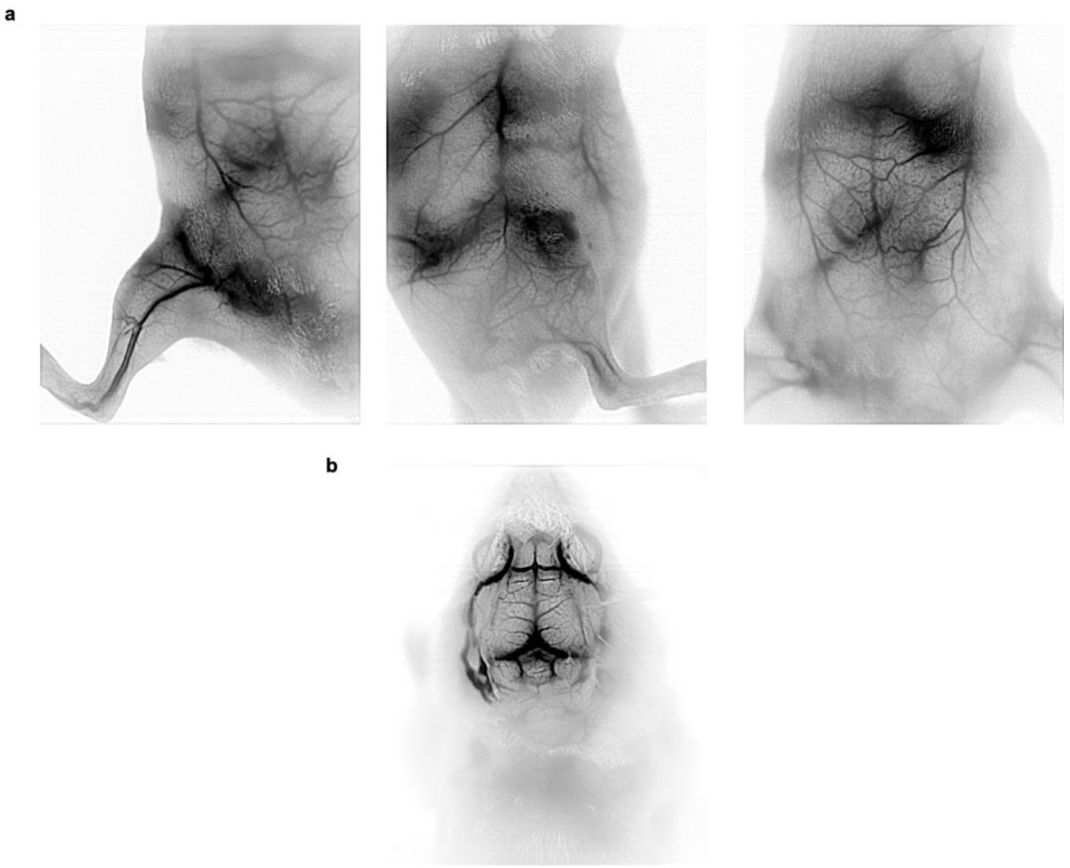

Figure S11. **a.** Noninvasive vascular imaging in vivo in mice, 15 min after i.v. injection of AuMHA/TDT (200 µL at 360 µM in PBS) with MCR imaging treatment ($\lambda_{exc.}$ 830 nm; LP1250 nm; 500 ms). **b.** Image of the brain (skin and skull removed) after MCR processing (reverse contrast). Reverse contrast provides a better visualization of small vessels.
